# Sensory information from a slipping object elicits a rapid and automatic shoulder response

**DOI:** 10.1101/724054

**Authors:** Carlos R. Hernandez-Castillo, Rodrigo S. Maeda, J. Andrew Pruszynski, Jörn Diedrichsen

**Author notes:** Correspondence to: Dr. Carlos R. Hernandez-Castillo, The Brain and Mind Institute, Western University, Western Interdisciplinary Research Building, Western University, Rm 4138, London, ON N6A 5B7, Canada.

## Abstract

Humans have the remarkable ability to hold, grasp, and manipulate objects. Previous work has reported rapid and coordinated reactions in hand and shoulder muscles in response to external perturbations to the arm during object manipulation; however, little is known about how somatosensory feedback of an object slipping in the hand influences responses of the arm. We built a hand-held device to stimulate the sensation of slipping at all five fingertips. The device was integrated into an exoskeleton robot that supported it against gravity. The setup allowed us to decouple somatosensory stimulation in the fingers from forces applied to the arm— two variables that are highly interdependent in real-world scenarios. Fourteen participants performed three experiments in which we measured their arm feedback responses during slip stimulation. Slip stimulations were applied horizontally, in one of two directions, and participants were either instructed to follow the slip direction, or to move the arm in the opposite direction. Participants showed responses within ∼67 ms of slip onset when following the direction of slip, but significantly slower responses when instructed to move in the opposite direction. Arm responses were modulated by the speed but not the distance of the slip. Finally, when slip stimulation was combined with mechanical perturbations to the arm, we found that sensory information from the fingertips significantly modulated the shoulder feedback response. Overall, the results demonstrate the existence of a rapid feedback system that stabilizes hand-held objects.

**NEW & NOTHEWORTHY:** We tested whether the sensation of an object slipping from the fingers modulates shoulder feedback responses. We found rapid shoulder feedback responses when participants were instructed to follow the slip direction with the arm. Shoulder responses following mechanical joint perturbations were also potentiated when combined with slipping. These results demonstrate the existence of fast and automatic feedback responses in the arm in reaction to sensory input to the fingertips that maintain grip on hand-held objects.

## INTRODUCTION

Imagine that you are looking at your smartphone, while your partner is asking you a question. After you fail to respond to the question, your partner decides to get your attention by pulling your phone out from your hand. In this situation, your partner’s action would initiate a combined response of your upper limb and hand to stabilize your grasp and secure the device. How the nervous system rapidly uses haptic and proprioceptive feedback to appropriately respond in such complex real-world scenarios is an important question in sensorimotor neuroscience (Mazurek et al., 2018; Mathis et al., 2019).

Previous reports have shown evidence that the nervous system automatically increases grip force to prevent an object from falling when slip is detected (Cole and Abbs 1988; Jones and Hunter 1992; Johansson et al. 1996; Johansson and Westling 1984). In the case of self-initiated movements, these grip-force modulations are highly predictive (Danion and Sarlegna 2007; Diamond et al. 2015; Flanagan and Wing 1997; Hadjiosif and Smith 2015; Wolpert and Flanagan 2001). Within the arm, humans generate rapid and flexible motor responses in response to mechanical perturbations that compensate for the coupling between joints (for review see Pruszynski et al. 2012) and are modulated by task goals (Pruszynski et al. 2008; Pruszynski et al. 2016; Weiler et al., 2019).

Previous work has mainly characterized grip and upper limb responses independently— it is clear, however, that hand and arm responses need to be tightly coordinated for successful object manipulation (Smeets et al. 2019). To explore this coordination, Crevecoeur and colleagues (2016) applied loads to the arm joint while participants held and object in precision grip. Their results showed that hand muscles rapidly accounted for the perturbation direction in a goal-dependent manner. Thus, perturbation in the upper limb modulates grip force. In is unknown, however, whether there is a fast an automatic coupling between sensory information from the fingers (e.g., slipping object) and arm feedback responses.

To study how somatosensory information at the finger tips modulates arm responses, we designed a new device to emulate the sensation of an object slipping during grasping.

Importantly, the object slip could be manipulated independently from any loads applied to the shoulder or elbow joints. In real life, when somebody pulls an object you are holding, part of the force will be transmitted to your arm and sensed via the muscle spindles, resulting in a direct compensatory response of the arm muscles (Dimitriou 2014). Hence, any arm response in this scenario could be the result of proprioceptive information from the arm rather than from somatosensory information from the finger tips. To be able to disentangle these two sources of information, we mounted the device on a robotic exoskeleton, such that the forces inducing the slip sensation at the fingertips could be uncoupled from the forces applied to the arm. This allowed us to investigate the effect of the somatosensory information from the fingers, without the confounding influence of proprioceptive information at the arm.

We hypothesized that the sensation of an object slipping may trigger a rapid shoulder muscle response to compensate for the slipping direction. A priori, it was not clear whether such an automatic response would involve the arm following the direction of slip or opposing the direction of slip. In Experiment 1, we therefore compared responses under a “follow” or “against” instruction and found a much more rapid response when participants followed the direction of slip. In Experiment 2, we tested how the speed and distance of the slip would influence the rapid shoulder muscle response. Finally, Experiment 3 investigates how this mechanism interacts with mechanical perturbations applied to the shoulder joint, as occurs in real-world scenarios. This design allowed us to study somatosensory and proprioceptive perturbations in the hand and shoulder independently, as well as the interaction between them when these perturbations are combined.

## MATERIALS AND METHODS

### Participants

Fourteen human participants (aged 22.7 ± 3.7; 6 males, 8 females) with no known musculoskeletal or neurological diseases were invited to perform three experiments described below. Participants reported to be right-hand dominant and had normal or corrected-to-normal vision. The Office of Research Ethics at Western University approved all experimental procedures according to the Declaration of Helsinki, and all participants signed a consent form prior to participating in an experiment.

### Apparatus

Participants performed the experiments using a robotic exoskeleton (KINARM, BKIN Technologies, Kingston, Ontario, Canada) that permits flexion and extension movements of the shoulder and elbow joints in the horizontal plane intersecting the shoulder joint (Scott 1999). The KINARM robot can independently apply mechanical loads to the shoulder and/or elbow and record kinematic variables of these joints. Mechanical stimuli were delivered to the fingertips using a custom-built, computer-controlled stimulator box, designed to produce a slipping sensation at each of the five fingers (Figure 1a). The stimulator box was mounted to the KINARM (fixed in the hand plate) and participants grasped it during the task. The stimulator allowed position and speed control of the contact surfaces in one dimension for all fingers. The surface that contacted the fingertip was flat and had fine sandpaper (grit 800) as a surface finish. This stimulus surface was chosen to obtain sufficiently high friction between the contact surface and the skin without restraining the slider movement. The contact surface for each finger was 18 mm in the vertical plane and 40 mm in the horizontal plane. The range of movement of the sliders was 18 mm driven by high-speed digital servos (Power HD 3688HB; operation speed 0.07 sec/60°; stall torque 2.8 kg-cm). To measure the grip force of each individual finger, two load sensors (Honeywell FSG020WNPB) per finger were placed behind the sliders. Because the hand, arm, and the case of the finger-stimulation box were all fixed to the KINARM exoskeleton, slip stimuli delivered to the fingers did not induce any torque in the elbow or shoulder joints. The setup included an overhead screen and semitransparent mirror to show visual information. Each segment length of the robot was adjusted to fit the participant’s arm. Arm supports were selected according to the arm size and foam padding was used to reduce any undesirable arm movement. Throughout the experiment, direct vision of the entire arm and hand was occluded so that responses were guided only by somatosensory information.

**Figure 1:**
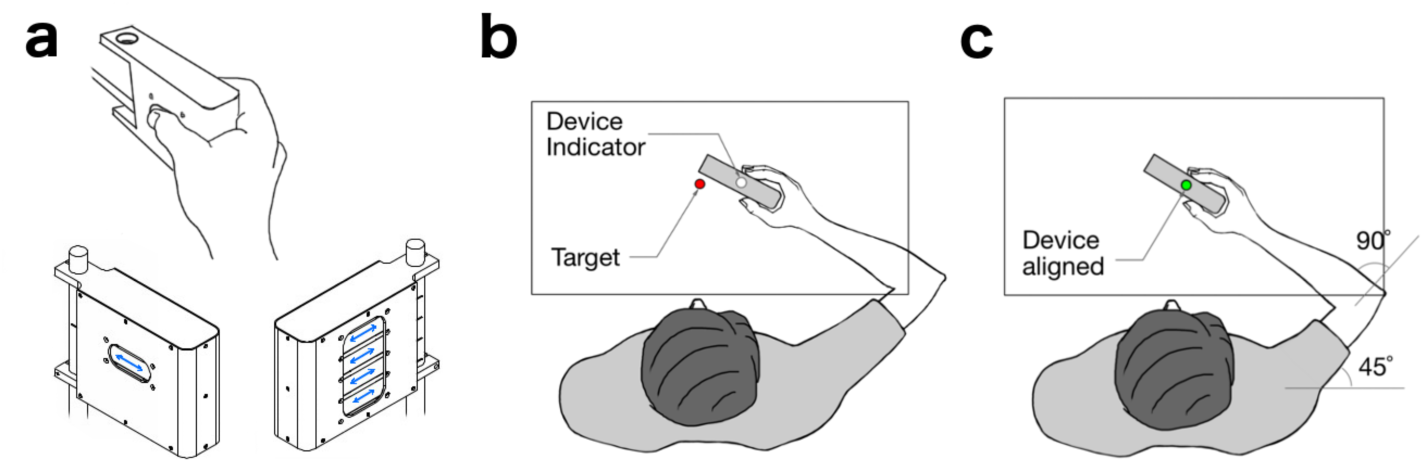
Stimulator device and experimental setup. (a) Left, right and holding view of the stimulator box. Blue arrows indicate the movement of the sliders. (b) In all experiments, participants held the stimulator box that could trigger a slipping sensation at the fingertips. Visual feedback of the device position (white circle) and the target position (red circle) were displayed in the same plane of motion. (c) Before each trial, participants were instructed to align the device visual feedback with the target feedback while accomplishing the baseline state conditions of position, grip force, and muscle pre-activation (see experimental paradigm). All visual feedback was then removed for the start of a trial (i.e., prior to the delivery of a mechanical slip, mechanical joint perturbation, or both).

### Experimental paradigm

#### Experiment 1

Rapid feedback responses. We hypothesized that the sensation of the object slipping in the finger tips would cause a rapid response in the arm. A priori we did not know whether this response would cause the arm to follow the object slip (to stabilize the object) or whether it would move the arm in the opposite direction (to resist the perturbation). We therefore designed a postural task in which the participants held the stimulator box while they felt the slip in one of two directions—either inward or outward with respect to the hand. In separate blocks, participants were either instructed to “follow the slip” or to “move against the slip”. If there exists a rapid and automatic coupling between slip sensation and arm response, the reaction in the “natural” direction should be substantially faster. The procedure began with the participant grasping the stimulator while seated in the exoskeleton. During all trials the direct visual feedback of the hand and arm was occluded, however, during the initial part of the experiment, a visual cursor (white circle: 1 cm diameter) indicating the position of hand was projected onto the mirror (Figure 1b). To start a trial, the participant had to fulfill three conditions: 1) Using visual feedback, participants had to align their hand (white cue) with the home target (red circle: 2 cm diameter) whose position corresponded to a shoulder angle of 45 degrees and an elbow angle of 90 degrees (Figure 1c). 2) After entering the home target, the exoskeleton gradually applied a background torque of 2 Nm to either the flexor or extensor muscles of the shoulder (arm pre-activation). Participants were instructed to keep their hand at the home target while grasping the stimulator. 3) Participants had to apply a grip force of 0.5 N ± 0.1 N between the thumb and the rest of the fingers. Once participants achieved these three conditions, all visual feedback was removed. Then, if participants maintained this baseline state for a random period between 250-500 ms (uniform distribution) the trial started. If participants failed to achieve/maintain this baseline state for 1 s the trial restarted from the beginning. For Experiment 1, participants were instructed to move their arm as fast as they could either in the same (to follow) or the opposite (go against) direction of the slip. To avoid any constraints on the movement, participants did not receive any instructions pertaining to the distance they should move. The slider displacement was 16 mm with a speed of 20 mm/s in either the inwards to outwards directions. Participants completed 240 trials in two blocks. Half of the participants received the instruction of “follow the slip” first and the other half received the instruction of “move against the slip” first. The order of slipping direction was randomized and participants completed 120 trials in each block. About 20 minutes were required to complete Experiment 1.

#### Experiment 2

Speed and distance of the slip. To test whether speed and distance of the slip could modulate the arm response, participants performed an accuracy task. We asked participants to precisely compensate for the slip of the sliders with an arm movement. Thus, if the participant felt that the sliders moved 1 cm in the forward direction within the device, the hand was required to also move 1 cm in the forward direction. We ask participants to move without delay from the slip onset. As in Experiment 1, a trial in Experiment 2 started when participants accomplished and maintained the baseline state. Mechanical slip occurred at one of two different distances and two speeds. Participants completed a total of 96 trials in this experiment. The instruction was to follow the direction of the slip as accurate as possible. The order of slipping distance (8/16 mm), velocity (10/20 mm/s), and direction (in/out) was randomized. About 20 minutes was required to complete Experiment 2.

#### Experiment 3

Combined slip and arm perturbations. In Experiment 3, we studied the interaction between simultaneous perturbations to the arm and slip stimulation at the fingertips. In this experiment, participants performed a postural task that required holding and keeping the stimulator box centered at a target. A mechanical load was applied at the shoulder joint, either alone or in combination with a slip stimulation to the fingers. The instructions to accomplish the baseline state were the same as in Experiments 1 and 2. At the moment of perturbation, the stimulator moved the sliders, and/or the KINARM robot applied a mechanical load at two different strengths (1 Nm or 2 Nm) at the shoulder joint. Participants were instructed to move the hand back to the original position (without visual feedback), as quickly as possible after perturbation onset. Participants completed a total of 96 trials in this experiment. The order of slip stimulation (present/absent) and strength of joint perturbation (1 Nm/2 Nm) was randomized. About 20 minutes were required to complete Experiment 3.

### Muscle activity

Surface EMG recordings were obtained from four upper-limb muscles involved in flexion or extension movements at the elbow and/or shoulder joints (pectoralis major clavicular head, PEC, shoulder flexor; posterior deltoid, PD, shoulder extensor; biceps brachii long head, BI, shoulder and elbow flexor and wrist supinator; triceps brachii lateral head, TRI, elbow extensor). Prior to electrode placement, the skin was cleaned and abraded with rubbing alcohol and the electrode contacts were covered with conductive gel. Electrodes (DE-2.1, Delsys, Boston, MA) were placed on the belly of the muscle, oriented along the muscle fiber, and the reference electrode (Dermatrode, American Imex, Irvine, CA) was attached to the clavicle. To assess the quality of each EMG signal, we performed a set of maneuvers known to elicit high levels of activation for each muscle in the horizontal plane. EMG signals were amplified (gain = 103) and band-pass filtered (20 – 450 Hz) by a commercially available system (Bagnoli, Delsys) then digitally sampled at 1,000 Hz.

### Data analysis

Data processing and statistical analyses were performed using MATLAB (The Mathworks, Natick, MA). All joint kinematics (i.e., hand position and joint angles) were sampled at 1000 Hz and then low-pass filtered (12 Hz, 2-pass, 4th-order Butterworth). EMG data were band-pass filtered (20-500 Hz, 2-pass, 2nd-order Butterworth) and full-wave rectified. EMG data were normalized to their own mean activity over the 200-ms period before slip perturbation onset when either shoulder flexor or extensor muscles were loaded by the exoskeleton (i.e., shoulder flexion or extension torque preload, 2Nm). All data were aligned on perturbation onset that could be either a mechanical slipping, mechanical joint perturbation, or both at the same time. To estimate the temporal onset of task related EMG activity for each participant, we used each participant’s EMG activity from two conditions to generate a time-series receiver operator characteristic (ROC) from 0 ms – 200 ms relative to perturbation onset. Briefly, ROC curves quantify the probability that an ideal observer could discriminate between two stimuli conditions: a value of 0.5 represents chance-level discrimination, whereas a value of 0 or 1 represents perfect discrimination (Green and Swets 1966). ROC curves were generated from the pectoral or deltoid muscle EMG activity, depending on the condition. We then fit the time-series ROC curves with a linear regression technique, which estimates the temporal onset of task-related EMG activity by determining when the time-series ROC curve diverges from chance-level discrimination (i.e., ∼0.5; see Weiler et al., 2015). We will refer to this time point as the divergence onset time.

Hand tangential velocity was used to determine the end of the hand trajectories. We performed different statistical tests such paired t-test and ANOVA when appropriate for each of the three experiments. Details of these procedures are provided below in the Results section.

Experimental results were considered statistically significant if the corrected p-value was less than 0.05.

## RESULTS

### Experiment 1: Automatic arm response in the direction of slip

In Experiment 1 participants were instructed to move the hand position via the shoulder joint as fast as possible either in the same (to follow) or in the opposite (go against) direction of the slip. If there exists a rapid and automatic coupling between slip sensation and arm response, the reaction in the “natural” direction should be substantially faster.

Figure 2a shows the task design, in which participants performed backwards or forward movements for the two slip directions (2×2 design, Figure 2e). The mean kinematics of the shoulder joint are shown in Figure 2 b and f for forward and backwards arm movement, respectively. For both arm movements, we found that following the slip (red traces) resulted in faster responses compared to moving against the slip (blue traces). The EMG data also revealed a faster ramping of agonist muscle activity when the participants followed the slip (Figure 2c,g). To quantify the difference in timing, first we estimated the onset of divergence from baseline activity for the two conditions (follow and against) in each participant. Indeed, for the forward arm movement (Figure 2c), participants performed faster responses when they moved in the same direction of the slip (mean onset time = 60.0 ms; SE = 0.2) compared to when they moved in the opposite direction (mean onset time = 148.1 ms; SE = 0.5). Then we calculate the divergence time between the two conditions for each arm movement. In both cases the divergence between In and OUT conditions was close to 67 ms (forward 67.1 ms SE 0.1 and backwards 67.1 ms SE 0.2). A paired t-test indicated a significant difference (t(13) = 2.11, p = 0.027). This behavior was similar for the backward arm movement (Figure 2g), showing a faster arm response when participants moved in the same direction of the slip (mean onset 78 ms) compared to when they moved to the opposite direction (mean onset time = 153 ms; t(13) = 2.37, p = 0.016).

**Figure 2.**
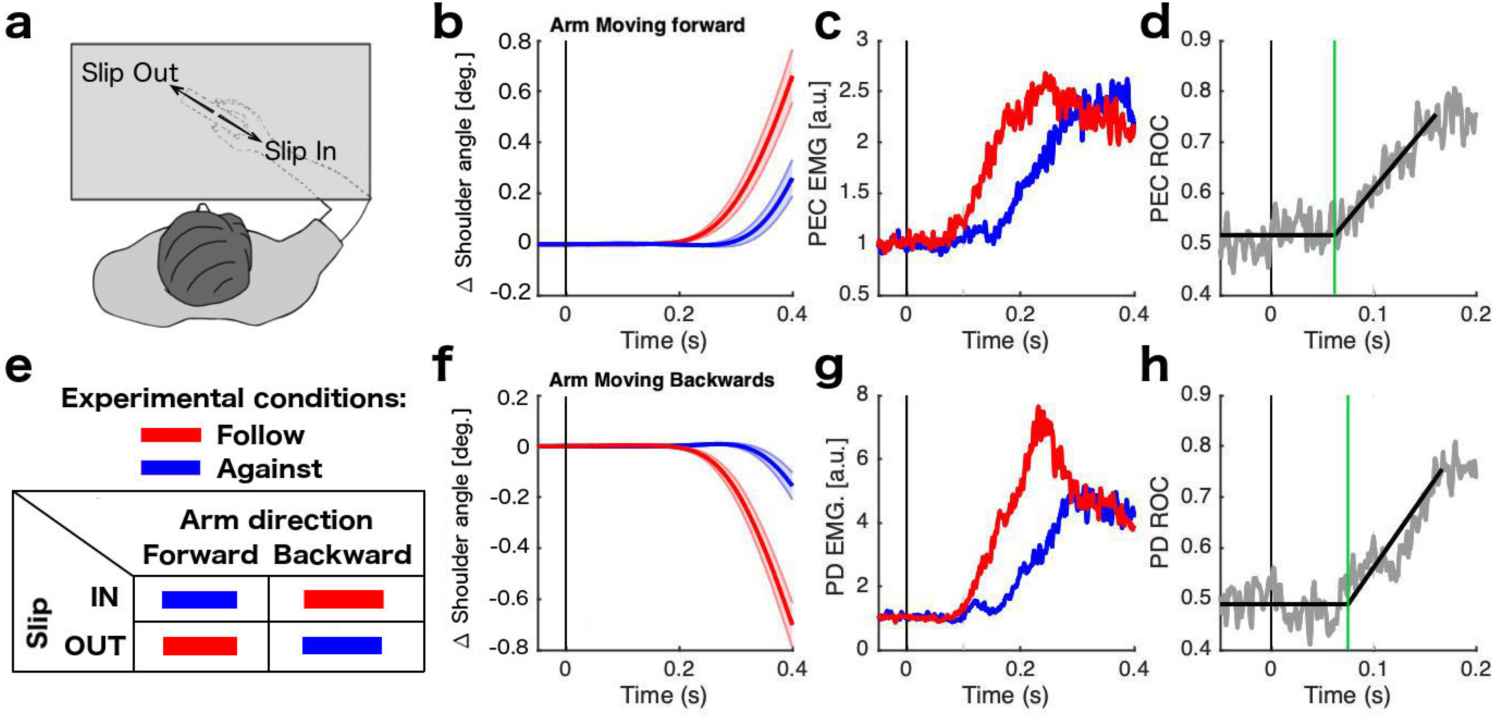
Shoulder responses related to slipping direction. During experiment 1, participants received slip stimulation in two directions (a) and they were instructed to move the arm either in the same (follow) or the opposite (against) direction of the slip (e). (b,f) shows the average kinematics of the shoulder joint. (c,g) Normalized muscle activity. (d,h) ROC curve of the divergence between follow and against conditions. (b,c,d) shows the results for a forward arm movement while (f,g,h) shows the results of backward arm movement. Shaded areas represent the standard error of the mean. ROC panels indicate in gray the ROC curve and in black the best fitted line. Green line indicates the timing of a significant difference of the muscle response for both conditions (red and blue). All Muscle activity traces correspond to the agonist shoulder muscle for each arm movement. deg. (degrees), a.u. (Arbitrary units). All data are aligned on slipping onset.

To investigate if the arm response to slip is different for forwards and backwards directions (shoulder flexion and extension), we determined the divergence onset time between the two conditions (follow and against) for each arm movement and then we performed a t-test between arm directions. This contrast did not reveal a significant difference (t(13) = 0.32, p = 0.374). Figure 2 d and h show time-series ROC curves from an exemplar participant fit with the linear regression technique that indicates the divergence onset time (green line) between follow and oppose movements in panels c and d, respectively.

These results show that the arm feedback response is faster when the arm movement is in the same direction of the slip as compared to when the participant moves in the opposite direction.

If there is an automatic response to follow the direction of a perceived slip, we would expect that some of the feedback responses under the “move against” instruction is produced in the wrong direction (i.e., in the direction of the slip). To test for this possibility, we carefully analyzed the paths of the hand during the trials. Figure 3a shows the average displacement trace of the hand position for each participant, showing that participants generally followed the instruction. However, on individual trials, participants made a number of errors. We defined an error as individual trials when the participant moved more than 1 mm away from the home position (either in the x or y axis) in a direction different from the correct quadrant (i.e., second quadrant for the forward movement, fourth quadrant for the backward movement; Figure 3b). Participants showed only a small number of errors when the arm movement followed the slip (3.1% of total trials) compared when the slip was opposite to the arm movement (26.9% of total trials). This difference was significant for both forward (t(13) = 3.59, p = 0.001) and backwards movements (t(13) = 3.21, p = 0.002).These results suggest that the response to follow a slipping object with the arm is not only fast, but also automatic—that is, it can intrude on a voluntary response and induce errors (Haith and Krakauer 2018).

**Figure 3.**
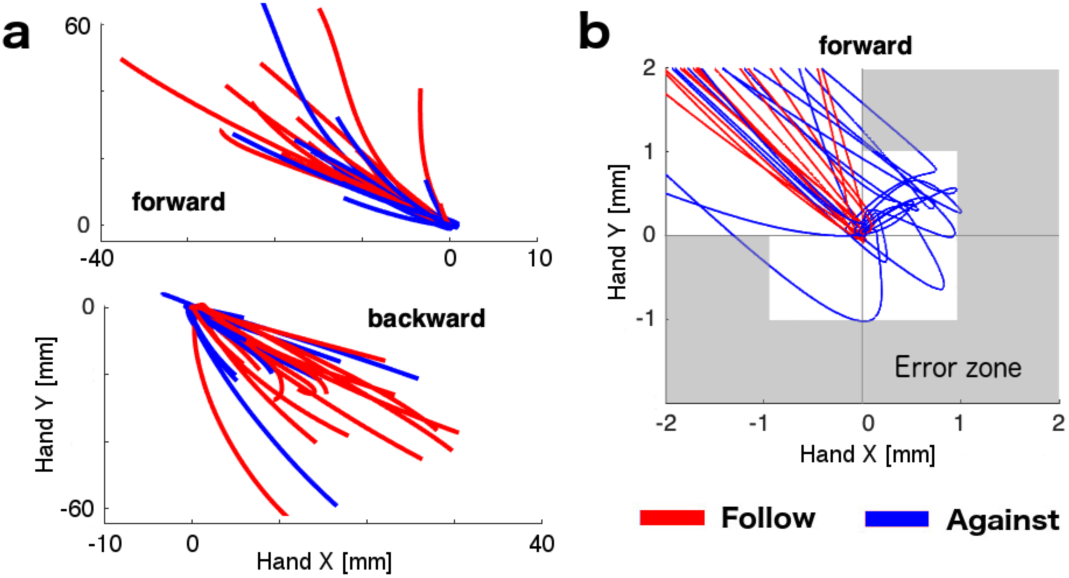
Hand paths for experiment 1. a) Each trace indicates the average path of each participant for both conditions—follow (red) and against (blue)—and both directions of arm movement (forward and backward). Paths start on the trial onset (at home position 0,0) and finish after 600 ms. b) Zoomed view of the home position in the forward movement. Gray area indicates the error zone for individual trials. Note that the image shows the average traces which hardly fall in the gray area, however individual trials marked as error trials exceed those limits.

### Experiment 2: Fast feedback responses vary with speed, but not with the distance of slip

In Experiment 1, we showed an automatic response of the arm that follows the slip sensation on the fingers. It has been shown that rapid responses can be modulated in a task-dependent manner to maintain limb stability (Shemmell et al. 2010). We therefore tested whether the characteristics of the slipping stimulus modulates the arm response, or if the arm responds equally to any slip sensation. We used two speeds and two distances for the slip stimuli (Figure 4a). To limit the overall number of conditions, we chose to study only forward arm movement with slipping in the direction out of the hand. Overall, we found that faster slips (orange colors in Figure 4c) elicited earlier (mean onset time = 67.0 ms, SE 0.6) muscle activity compared to slower slips (green colors; mean onset time = 114.3 ms, SE 0.7; t(13) = 3.99, p = 7.6e-4).

**Figure 4.**
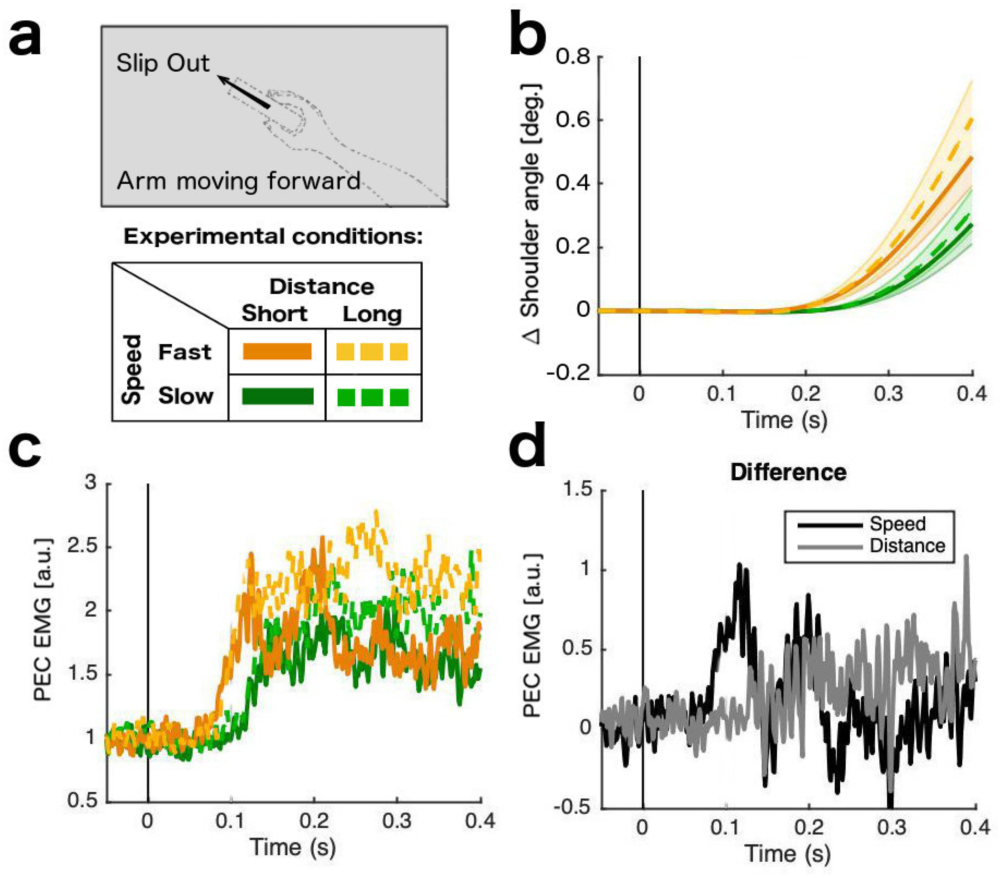
Shoulder responses according to different slip characteristics. During experiment 2, participants received slip stimulation in the “out” direction using two speeds and two distances (a) and they were instructed to move the arm following the slip. (b) Average kinematics of the shoulder joint. (c) Normalized muscle activity. (d) Gray line shows the difference between Fast Long and Fast Short (Distance) while black line shows the difference between Fast long and Slow long (Speed). All Muscle activity traces correspond to the agonist shoulder muscle for each arm movement.

However, the muscle activities resulting from the two slip distances using the same slip speed (solid vs dashed lines of the same tone), were not significantly different for either slow slip (t(13) = 0.89, p = 0.194) or fast slip (t(13) = 1.36, p = 0.097). These results suggest that the speed of the slipping has a stronger effect on the early arm response, as compared to slip distance (Figure 4d).

The explicit task goal in Experiment 2 was to move the hand the same distance as the sensed slip (i.e., the displacement of the device sliders). Although participants’ movements did not exactly match the distance (8 or 16 mm), the average displacement showed a clear effect of the slip characteristics on the final position of the participant’s hand (Figure 5a). The slip distance (short vs. long) showed a clear influence on the final position, both in the slow (Figure 5b, t(13) = 5.40, p = 1.2e-4) and fast conditions (t(13) = 4.37, p = 7.5e-4). Although the instructions emphasized an accurate compensation for the slip distance, the speed of slip also had a significant influence on hand displacement for both the short (t(13) = 1.83, p = 0.044) and long slips (t(13) = 2.19, p = 0.023). An ANOVA also showed a significant interaction between slip speed and distance (F(3,39) = 20.3, p = 5.6e-6), resulting from a larger influence of speed in the long distance condition as compared to the short distance condition.

**Figure 5.**
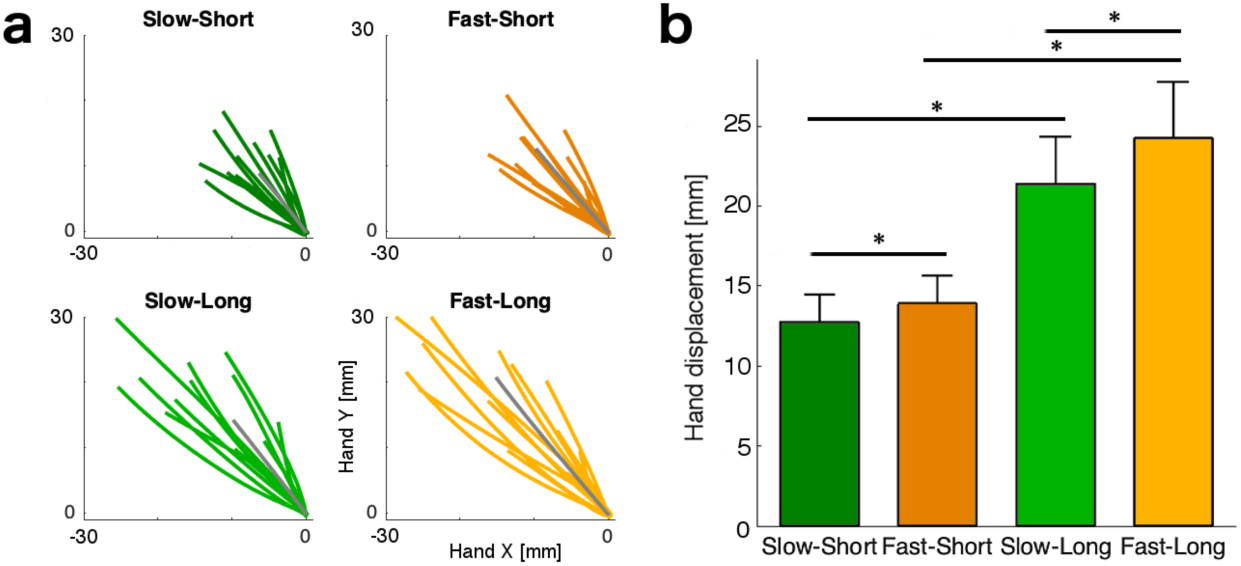
Hand paths for experiment 2. a) Each color line indicates the average path of each participant for each condition for the forward arm movement. Gray line indicates the mean path of the group. Paths start on the trial onset (at home position 0,0) and finish when the participant stops movement (tangential velocity < 30% of the maximum velocity of each trial). b) Average hand displacement from the home target to the end of the movement for each condition.

Overall, these results show that the initial arm response is mostly dictated by the speed of the slip. In contrast, the overall response of the arm took into account the displacement of the slip to achieve the behavioral goal, but still was slightly biased by the initial speed.

### Experiment 3: Slip modulates response to arm perturbation

In real-world scenarios our nervous system needs to integrate information from the finger tips with information from the arm to optimally resist perturbations delivered to a hand-held object. Our setup uncoupled these sources of information between the hand and the arm, allowing us to observe the effect of slip stimulation in isolation. But how do feedback from the hand and arm interact when perturbations occur simultaneously with slip stimulation? It is possible that the local arm feedback loop completely overwrites any modulation from the sensation from the fingertips. Alternatively, the two sources of information may be combined in the final response. In Experiment 3, we investigated whether the slip sensation at the fingers modulates the arm’s response to a slipping object during an external arm perturbation (either 1Nm or 2Nm). We asked participants to bring the object back to the home position as fast as they could after the perturbation. Figure 6a shows the task setup and Figure 6b the response of the arm to an external mechanical shoulder extension perturbation alone (dashed lines), and to an external perturbation plus slipping in the opposite direction (i.e., out of the hand; solid lines).

**Figure 6.**
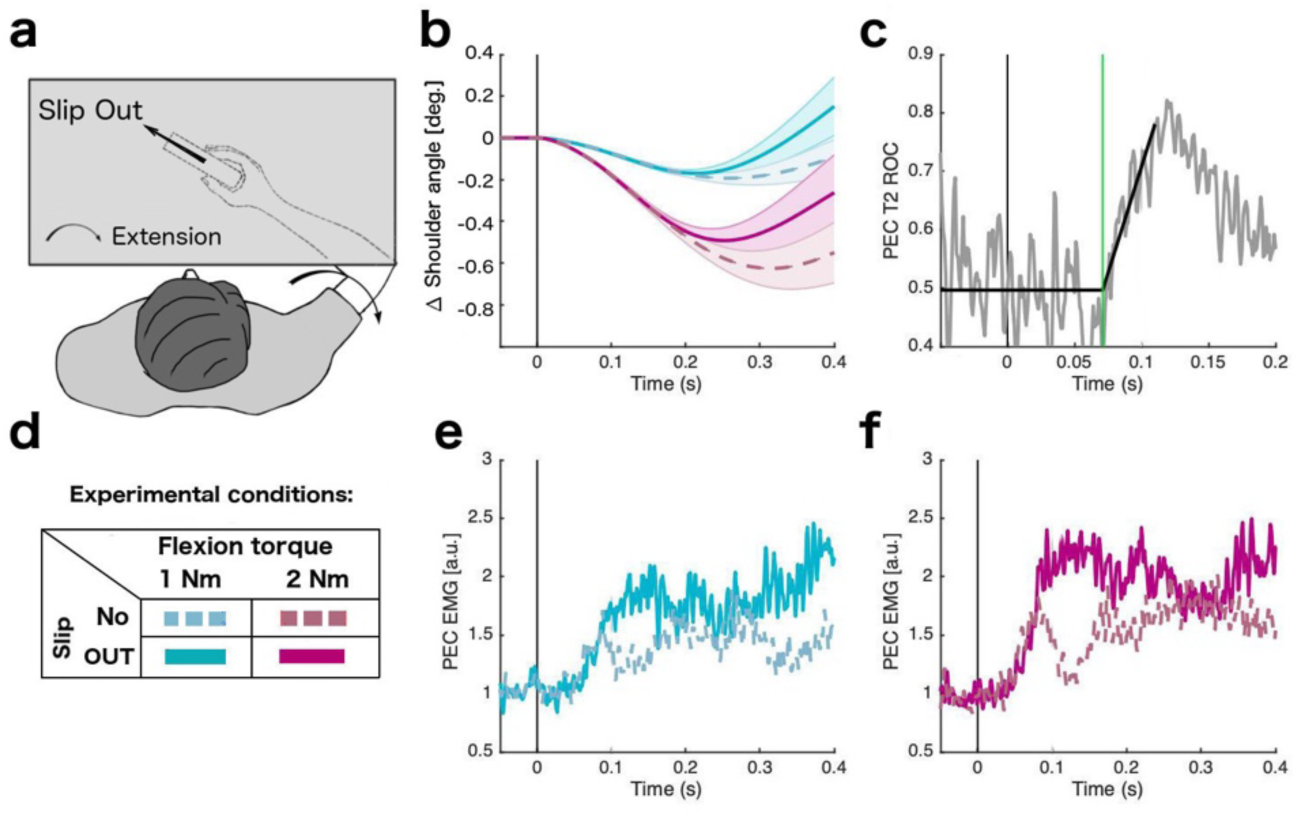
Arm responses related to combined torque and slip. During experiment 3, participants received either a flexion torque (1 Nm or 2 Nm) or a flexion torque plus slip stimulation (out direction) (a,d). Participants were instructed to move the stimulator cursor back to the original position (without visual feedback). (b) Average kinematics of the shoulder joint. (e,f) Normalized muscle activity. (c) ROC curve of the two conditions (torque and torque plus slip) using 2 Nm. ROC panel indicate in gray the ROC curve and in black the best fitted line. Green line indicates the timing of a significant difference of the muscle response for both conditions. All Muscle activity traces correspond to the agonist shoulder muscle for each arm movement.

As expected, the 2 Nm torque produced larger arm displacements than the 1 Nm perturbation (Figure 6b). For both perturbation levels, however, the position of the arm moved back to the original position faster when the slip was included in the perturbation, as compared to when it was absent (torque alone). Although the onset of the EMG activity did not change significantly, the EMG signal showed a significantly higher activity when the slipping stimulation was present (Figure 6e, f). To determine the onset of this modulation, we computed the area under the ROC curve for each time point and determined the divergence between trials with and without slip present using linear regression (see methods). The mean onset time for 1 Nm was 98.2 ms, SE 0.9; while for 2 Nm we found a mean onset time of 71.3 ms, SE 0.9 (Figure 6c). For both torques the EMG signal was significantly higher when the slip was present immediately after the divergence time: 1Nm (t(13) = 2.95, p = 0.005) and 2 Nm (t(13) = 5.27, p = 8.0e-5). This result suggests that the direct perturbation in the arm does not override the slip sensation from the fingertips, but that both are integrated to produce a combined feedback response.

Participants were relatively accurate in returning to the home target when they received a mechanical torque in the arm. Figure 7a shows the average hand path of each participant for each condition. As expected, the stronger perturbation (2 Nm) resulted in higher variability in the end position of the hand, but overall, participants stopped close to the home position. When the slip was present, however, participants tended to overshoot, ending the movement farther away from the home position compared to the respective control (torque alone). An ANOVA comparing the individual end positions showed a significant main effect of the slip (F(3.39) = 13.8, p = 1.6e-5). We also found a significant interaction between torque and slip (F(3,39) = 11.4, p = 2.3e-6) - the difference between the control and combined condition (torque plus slip) was higher for the 1 Nm perturbation (t(13) = 5.38, p = 1.2e-4) than for the 2 Nm perturbation (t(13) = 2.73, p = 0.017) (Figure 7b). Overall, slip information biased participants to respond more strongly to the perturbation, ultimately leading to a less accurate performance.

**Figure 7.**
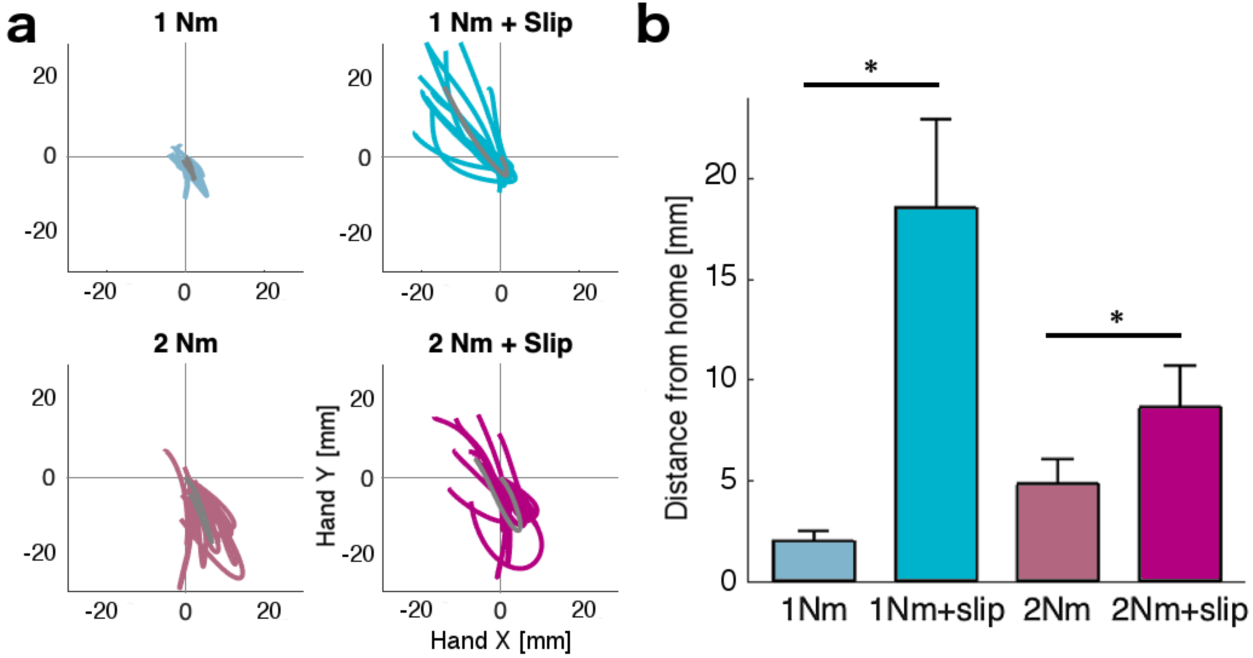
Hand paths for experiment 3. a) Each color line indicates the average path of each participant for each condition. Gray line indicates the mean path of the group. Paths start on the trial onset (at home position 0,0) and finish when the participant stopped the movement (tangential velocity < 30% of the maximum velocity of each trial). b) average hand displacement from the home target to the end of the movement for each condition.

## DISCUSSION

Taken together, our results establish the existence of a fast and automatic arm response that follows the direction of an object slipping from the hand. We were able to reveal this response by artificially uncoupling the slip sensation on the fingertips from the forces acting on the shoulder joint, two variables that are often coupled in real-world situations. In our experiment, the stimulator device was fixed to the robot structure and the hand and arm of the participant were secured with foam padding to prevent any undesired movement within the device. Thus, the slip stimulation did not produce a torque to the arm and the torque applied to the arm did not cause slip of the device, allowing us to assess the arm responses associated with the slipping sensation alone. We report three principal findings. First, we found a fast and automatic feedback response in shoulder muscles when following the direction of a slip stimulus at the fingertip with an onset latency of ∼67 ms. Second, this rapid feedback response of the shoulder muscles was modulated by the speed but not by the distance of the slip. Third, responses to mechanical perturbations applied to the upper limb were potentiated when combined with object slip in the direction opposite to the perturbation.

### Automatic response following a slipping object

Previous work has long demonstrated that the sensation of slip at the fingertips can trigger very rapid increases in grip force (Delhaye et al., 2014; Häger-Ross et al., 1996; Häger-Ross and Johansson, 1996; Crevecoeur et al., 2017; Cole and Abbs 1988; Häger-Ross et al., 1996; Jones and Hunter 1992; Johansson and Westling 1984). Here we found that slip at the fingers also induces a rapid and automatic shoulder muscle response that moves the arm in the direction of the slip. This automatic response was revealed by instructing participants to either follow the slipping direction or to move against it—a paradigm similar to anti-saccade or anti-reach approach (Munoz and Everling, 2004; Gail and Andersen, 2006; Day and Lyon, 2000). Specifically, we found substantially faster responses when the participants were instructed to move their arms in the same direction of the slip as compared to when instructed to move in the opposite direction. If the responses had been arbitrary and fully deliberate, both instructions should have led to the same latency.

A related observation comes from a bimanual haptic tracking task (Rosenbaum et al., 2006). In this study, participants were instructed to follow a moving object using the tactile information from the fingertip that made contact with the object. The results show that participants could follow two independent spatial trajectories with their two hands without interference—something that is very hard to achieve during voluntary movements (Kennerley et al., 2002). The lack of interference clearly argues for the existence of an automatic response that guides the arm in the direction of a perceived slip.

What is the functional relevance of this automatic response? It is most likely that is serves to facilitate stability of a hand-held object. When an object slips from our grasp, it is essential to follow the movement of the object with the arm to prevent the object from completely slipping from our grasp. Even smaller movements of the object within the grasp should be prevented, as the finger grasp positions are chosen to balance the object in the hand to avoid object rotation (Mackenzie and Iberall 1994).

Consistent with a functional role in object stabilization, we showed in Experiment 2 that the arm responses scale with the initial speed of the slip. For grip force increases, such modulation has been well demonstrated (Häger-Ross and Johansson, 1996; Cole and Abbs, 1988; Crevecoeur et al., 2017). In contrast, we found no modulation in the initial shoulder muscle responses when the grasped object slipped at two distinct distances. This was expected, as at the onset of slipping in either condition (short or long distance), the same somatosensory information was transmitted to the nervous system. The differences between the two distances would therefore only become available when the short distance perturbation was completed. Indeed, the later responses and hand distance traces were clearly influenced by the length of the slip. These results provide evidence that the automatic response takes into account afferent feedback from the digits in an adaptive, time-sensitive, and appropriate manner but the contribution of tactile and or muscle afferent feedback remains to be elucidated. The muscle activity latency of the following response of the arm (∼67 ms) indicates that the response can be produced faster than normal voluntary responses, which usually have a time scale of 100-150ms. Similar latencies have been reported in previous work for other automatic responses, including the increase in grip force following a load perturbation in the fingertip (Cole and Abbs 1988; Crevecoeur et al. 2017), or a perturbation to the upper limb (Crevecoeur et al. 2016). The short latency indicates that these responses are not generated by the normal polysynaptic cortical circuits that underlie voluntary and potentially arbitrary responses. The ∼67ms response also suggests that these automatic responses are not generated exclusively at the level of the spinal cord, as known spinal reflexes (i.e., to muscle stretch) occur within ∼20-50ms (Weiler et al., 2019; Pierrot-Deseilligny et al., 2012). Feedback responses following mechanical perturbations that arise >50 ms can potentially engage spinal, subcortical, and cortical areas (Cheney and Fetz 1984; Evarts and Tanji, 1976; Pruszynski et al. 2011; Pruszynski et al. 2014; Omrani et al., 2016; for review see, Scott, 2016). While the neuroanatomical substrate that underlies these automatic responses remains to be determined, our study predicts that somewhere in the nervous system, neurons that project to shoulder muscles must receive relatively direct sensory input from tactile sensors in the hand. The response we describe here is similar to the nociceptive withdrawal reflex, where cutaneous inputs drive muscle responses to move the body away from a potentially dangerous stimulus (Sherrington, 1910). Indeed, careful mapping of the withdrawal reflex has revealed an intricate relationship between the location of the nociceptive stimulus and which muscles are recruited to best move the limb away from the stimulus (Schouenborg and Kalliomäki, 1990; Levinsson et al., 1999). A similar mapping and neural substrate could potentially underlie the responses observed here. It should be noted, however, that the direction of function of the following response is substantially different from the withdrawal reflex and thus may require different descending modulation and/or directly engage brainstem and cortical circuits also known to receive rapid somatosensory inputs (Scott, 2016).

### Combination of slip information with local muscle stretch

In our experimental setup, we artificially dissociated the slip information and the torques acting on the arm. In real world scenarios, however, a perturbation to a hand-held object will induce both slip of the object in the hand and a torque at the shoulder joint. In other possible scenario, the salience of the torque in the shoulder joint (proximal proprioceptive) will be higher in comparison to the stimulation on the fingertips (distal somatosensory) resulting in a preponderant response to the local perturbation in the joint. If the automatic response revealed in the first two experiments indeed functions to stabilize the hand-held object, it must also be functional in combination with stretch to the shoulder joint itself. The results from Experiment 3 clearly show that the automatic response to a slip is not overridden by the presence of a perturbation to the shoulder, but rather combines with this locally generated response.

The experimental situation corresponds to the natural scenario in which a perturbation to the arm causes a sudden acceleration of the limb. The inertia of the object then induces a slip of the object in the opposite direction. If such slip is detected, the resistive reaction of the arm is amplified, stabilizing the grasp on the object. While not reported here, pilot experiments also indicated that this amplification was not observed when the object slip was in the same direction of the arm perturbation. This arises from forces that are applied directly to the object, in which case the arm should be more compliant to maintain a stable object grasp.

Processing of sensory information from the hand and the upper limb have been largely studied in isolation (Delhaye et al. 2018; Scott, 2016); however, the integration of these two sources of information for limb control suggest a confluence of these sensory sources on motor structures. For example, spinal, subcortical (i.e., thalamus), and cortical (i.e., somatosensory cortex) structures are known to receive information from both tactile sensors and muscle spindles (Delhaye et al. 2018; Scott, 2016; Kim et al, 2015; Picard and Smith, 1992). Despite that our experiment did not provide data to test a specific way of integration, one possibility is that the observed combination might take place in regions that receive both types of information. Alternatively, it remains possible that the signals are processed separately, and the combination arises during convergence onto spinal motor neurons.

One limitation of our experiments is that we could only study a limited set of slip directions in the horizontal plane. However, if the function of this automatic response is to stabilize hand-held objects, the arm’s response to slip should adapt flexibly to the configuration of the arm in space, and to the configuration of the object in the hand. This would imply that slip at the fingertips can also modulate automatic responses around the elbow joint. Such flexibility remains to be experimentally shown. Other limitation of our setup is that regardless that we try our best to constrain the arm and hand movement in the exoskeleton, it is impossible to completely suppress any small change in finger configuration, and as a consequence afferent feedback from the finger muscles was also likely contributing to some extent.

In summary, our paper demonstrates that somatosensory information at the hand elicits rapid motor corrections in the shoulder that are suitable to stabilize hand-held objects, are sensitive to the slipping direction and speed, and are integrated with local reflex responses at the shoulder.

## Acknowledgements

This work was supported by a Scholar award from the James S. McDonnell foundation, and a discovery grant from the National Science and Engineering Research Council of Canada (RGPIN-2016-04890, both to JD). Additional support came from the Canada First Research Excellence Fund (BrainsCAN). C.H.C. received a postdoctoral fellowship form the Brain and Mind Institute. R.S.M. received a salary award from CNPq/Brazil. J.A.P. received a salary award from the Canada Research Chairs program.

